# Foveal feedback supports peripheral perception of both object color and form

**DOI:** 10.1101/690305

**Authors:** Kimberly B. Weldon, Alexandra Woolgar, Anina N. Rich, Mark A. Williams

**Affiliations:** Center for Magnetic Resonance Research, University of Minnesota, Minneapolis, MN, USA; Perception in Action Research Centre (PARC), Department of Cognitive Science, Faculty of Human Sciences, Macquarie University, Sydney, NSW, Australia; ARC Centre of Excellence in Cognition and its Disorders, Macquarie University, Sydney, NSW, Australia; Medical Research Council (UK) Cognition and Brain Sciences Unit, University of Cambridge, Cambridge, UK

**Author notes:** Corresponding author (KBW).

## Abstract

Evidence from neuroimaging and brain stimulation studies suggest that visual information about objects in the periphery is fed back to foveal retinotopic cortex in a separate representation that is essential for peripheral perception. The characteristics of this phenomenon has important theoretical implications for the role fovea-specific feedback might play in perception. In this work, we employed a recently developed behavioral paradigm to explore whether late disruption to central visual space impaired perception of color. First, participants performed a shape discrimination task on colored novel objects in the periphery while fixating centrally. Consistent with the results from previous work, a visual distractor presented at fixation ~100ms after presentation of the peripheral stimuli impaired sensitivity to differences in peripheral shapes more than a visual distractor presented at other stimulus onset asynchronies. In a second experiment, participants performed a color discrimination task on the same colored objects. In a third experiment, we further tested for the foveal distractor effect with stimuli restricted to a low-level feature by using homogenous color patches. These two latter experiments resulted in a similar pattern of behavior: a central distractor presented at the critical stimulus onset asynchrony impaired sensitivity to peripheral color differences, but, importantly, the magnitude of the effect depended on whether peripheral objects contained complex shape information. These results taken together suggest that feedback to the foveal confluence is a component of visual processing supporting perception of both object form and color.

## Introduction

Visual object recognition is traditionally thought to conform to a bottom-up, feedforward model of processing that begins with the extraction of low-level object information in early visual areas [1,2]. From there, visual information proceeds along a hierarchy of cortical regions representing increasingly complex information. In addition, feedback connections from higher to lower visual areas also have an important role in visual perception, such that feedback modulates or attunes feedforward information [3–5]. Williams et al. [6] used multi-voxel pattern analysis of fMRI data to demonstrate that information about the category of novel objects [7] presented in the observer’s periphery could be decoded in cortical regions that corresponded to central, foveal visual space, an area far removed from the stimulus input. The authors attributed this to a feedback process, as the fovea remained unstimulated throughout the experiment. The results from Williams et al. [6] suggested a new type of feedback mechanism - one that is capable of constructing a new and separate representation of peripheral object information. Critically, stronger representation of peripheral object category in foveal retinotopic cortex correlated with better behavioral performance on the task, implying an important role for this representation in perception.

A follow-up transcranial magnetic stimulation (TMS) study by Chambers, Allen, Maizey, and Williams [8] showed that integrity of the foveal region at a timeframe consistent with feedback is essential for peripheral perception. In that study, observers performed a task similar to that in Williams et al. [6]. Observers fixated centrally while discriminating between novel objects that briefly appeared in the observer’s periphery. A TMS pulse applied to the occipital pole selectively impaired perceptual discrimination sensitivity of peripheral objects when applied ~350ms *after* stimulus onset compared to a TMS pulse applied at other points in the course of a trial. TMS applied at stimulus onset asynchronies (SOAs) from 150ms prior to stimulus onset to 250ms post-stimulus onset, as well as beyond 400ms post-stimulus onset, did not have the same disruptive effect on discrimination sensitivity. Taken together, these studies suggest a form of feedback that constructs a representation of objects removed from the associated visual input and, further, that this feedback is behaviorally relevant.

To date, studies examining the foveal feedback phenomenon have largely employed a relatively difficult behavioral task where the participants discriminate between briefly-presented novel greyscale objects [9,10]. However, in Williams et al. [6], the authors included one experiment where participants performed a color discrimination task on colored objects presented in the periphery. In that experiment, unlike the shape discrimination task, the authors did not find information about object form at the fovea, raising the possibility that foveal feedback is related to the task at hand. The authors did not, however, test whether color information could be decoded in foveal retinotopic cortex. Therefore, it is unknown whether foveal feedback is limited to carrying general shape information of visual stimuli, or if it may function for any one object characteristic related to the task being performed.

We have previously reported a behavioral measure of foveal feedback [10]. In brief, participants perform a discrimination task on achromatic novel objects briefly presented (~100ms) in their periphery while fixating centrally. An achromatic visual distractor presented at fixation impairs discrimination sensitivity when it appears 117ms after target onset, after the targets have disappeared from the display. This disruption in discrimination sensitivity at +117ms post-stimulus onset reliably occurs when a central distractor is presented to the observer at a time entirely disparate from the target presentation, and is more pronounced compared to distractor onsets at other stimulus onset asynchronies (SOAs), including SOAs later in a trial (e.g., more than 250ms). In a previous paper, we termed this temporally-specific disruption of peripheral discrimination sensitivity the “foveal distractor effect”. We [10] also demonstrated spatial specificity of this effect: discrimination sensitivity was not similarly impaired when a visual distractor was presented in the periphery at the critical SOA. This behavioral paradigm demonstrates the spatial and temporal specificity of foveal feedback and is an efficient method for investigating how feedback influences peripheral perception (see also 6, 7).

In the present set of experiments, we used the paradigm described in [10] to test whether the foveal distractor effect is specific to perceptual discrimination between object shapes, or if it also occurs during tasks requiring discrimination of another object characteristic, in this case, color. Color is a useful characteristic to use with these stimuli as its manipulation does not interfere with the fine spatial details of the novel objects. Further, it is unknown whether color information about peripheral objects can be decoded from foveal retinotopic cortex during a color discrimination task [6], but, in light of evidence from electrophysiology research in monkeys suggesting cortical layers receiving feedback connections from higher visual areas may be selective for chromatic information [12], such an outcome is feasible. If feedback to foveal retinotopic cortex contains behaviorally-relevant information about peripheral objects, then disruption to foveal visual space should disrupt discrimination sensitivity of objects in the viewer’s periphery when they perform both shape and color-discrimination tasks. In Experiment 1 we replicate the foveal distractor effect using colored novel objects (as opposed to achromatic objects) in a task where participants discriminate between the objects’ shapes, while ignoring their colors. A central distractor presented 117ms after the onset of the targets impaired discrimination sensitivity of object shape in the periphery compared to distractors presented at SOAs very early or later in the trial. In Experiment 2 we used the same stimuli used in Experiment 1 but altered the task: participants were required to discriminate between the target colors while ignoring their shapes. To pre-empt our results, a visual distractor presented at fixation impaired peripheral discrimination sensitivity of color in the periphery, again only at the critical SOA.

Research on the cortical processing of color suggests that the neural computations related to form and color are strongly linked in early visual areas [for a review, see 13]. Early coupling of chromatic signals with other visual object characteristics such as orientation [14–16] and figure-ground segregation [17] have been well documented. Multi-voxel pattern analysis of fMRI data shows that object representation in early visual cortex does include information about the conjunction of color and object shape information [18]. Taking this into account, it is unclear, based on the results from Experiment 2, whether the foveal distractor effect is occurring due to the disruption of task-relevant color information in and of itself, or if the effect is occurring as a result of bound color information to complex object form. We addressed this question in Experiment 3 by removing complex shape information from the targets and requiring participants to discriminate between circular patches restricted to low-level color information. Our results indicate that the disruption of peripheral color discrimination sensitivity in the absence of complex shape information remains temporally-specific; on the other hand, the strength of that disruption is flexible and task-dependent.

## Experiment 1: Discriminating Form

### Materials and methods

#### Participants

Twenty participants, screened for normal or corrected-to-normal visual acuity as well as normal color vision using Ishihara color plates, were recruited for Experiment 1. One participant’s data were not used in the analysis due to chance-level performance, leaving the datasets of 19 participants (15 female, 4 male; mean age = 23.3 ± 4.55 years) for analysis. Participants received either course credit or $15 for their participation and gave informed consent. All experiments in this study were approved by the Macquarie University Human Research Ethics Committee.

#### Stimuli and apparatus

Sixteen stimuli were selected from a set of 1296 pre-generated “smoothie” stimuli [7]. These 16 exemplars were selected to represent the most extreme variations in the larger set. Using Matlab (Mathworks), each of the 16 exemplars was covered with a colored, transparent mask created in CIE L*c*h color space. Every colored mask had a luminance value of 85 and a chroma value of 38. The colored masks varied in hue angle from 0**°** (red) to 200**°** (blue) in steps of five degrees, resulting in a full stimulus set of 656 objects. We used a large range of colors to mimic the variability in the shapes of the exemplars. A further smoothie stimulus, which was not one of the 16 main exemplars, was selected for use as a visual distractor. This distractor was covered with a colored mask that had a hue value of 63, which was not one of the possible target colors. In this way, it was possible for the distractor object to vary in degree of similarity, to the color and/or shape of either target while never being identical to either characteristic. Each stimulus subtended ~1.5**°** of visual angle.

Experimental sessions took place in a dimly-lit, windowless laboratory at Macquarie University, Sydney. Stimuli were presented on an sRGB-calibrated 27in Samsung SyncMaster AS950 monitor at a resolution of 1920×1080 pixels and a refresh rate of 120Hz. We tracked fixation of the right eye with an Eyelink 1000 remote eye-tracker at 500Hz. The camera and infrared illuminator were mounted in front of the participant below the desktop display so that the screen was not obscured.

#### Training procedure

Prior to the experiment, participants were trained on a basic discrimination task (with no central distractor). First, a white fixation cross was displayed for 315ms. Then, two colored target objects were displayed for 417ms in the upper left and lower right quadrants of the screen. The targets were presented in these same locations throughout the training tasks and the experiments (Fig. 1). Participants were instructed to maintain fixation on the central cross throughout each trial and determine if the two targets in the array were different shapes or if they were identical in shape as quickly and accurately as possible, while ignoring the color of the targets. In “same” trials, the targets were always presented in the same orientation. In half of the trials, these target stimuli were different shapes, chosen at random from the larger set of 16, and in the other half they were identical shapes. The targets, regardless of whether they were the same or different shapes, always differed in color by a hue angle of 60°. The degree of color difference was selected based on pilot data, such that participants’ performance on a shape discrimination task would be similar to their performance in a color discrimination task using the same stimuli (see Experiment 2). Participants had 2000ms to respond with their right index finger or middle finger on the keyboard to indicate a “same” or “different” judgment, respectively. Following each response, participants were given onscreen accuracy feedback. After a 2000ms interstimulus interval, the next trial commenced automatically. Trials where the participant’s eye gaze drifted more than 2° from the center of the display were coded as incorrect during training.

**Fig 1.**
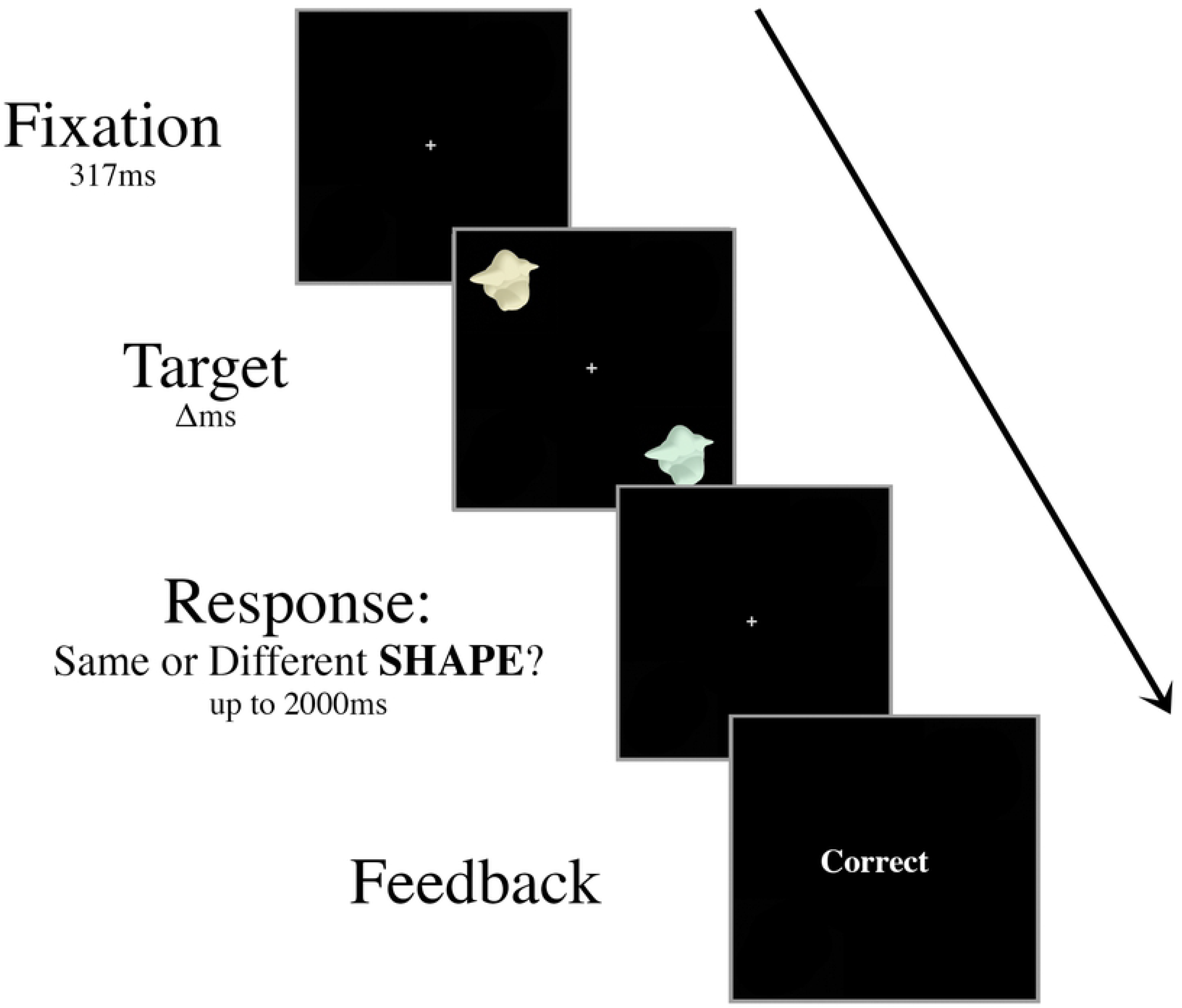
Schematic of an example “same” trial in the Experiment 1 training task. Targets were presented for decreasing durations (Δ: 417ms, 267ms, then 117ms) during training. Participants were instructed to ignore the color of the targets and judge only if the shapes of the targets are identical. In this example, the two targets are different colors but the same shape, requiring a “same” response. Training continued until the participant was able to perform above 70% accuracy with a 117ms presentation time across a single block of ten trials.

Trials were presented in blocks of ten. Once participants could perform the discrimination task with >70% accuracy across a single block with a target display duration of 417ms, the presentation time of the targets decreased to 267ms. Participants repeated the training procedure until they were able to perform the task with >70% accuracy in a block. Then, the presentation time of the targets further decreased to 117ms, which reflected the timing conditions in the experiment. Training continued until participants were able to make at least 70% correct discriminations when the target array was displayed for 117ms, while maintaining fixation throughout the block. In general, participants were able to complete the training within 20 minutes.

#### Experimental procedure

The procedure for Experiment 1 was similar to the training procedure with two major changes: there was a fixed target presentation duration of 117ms and a distractor object appeared at fixation once during each trial (Fig. 2). At the beginning of each trial, a white central cross was displayed for 567ms. In each target display, two colored targets were displayed for 117ms in opposite diagonal locations (upper left and lower right-hand quadrants of the screen), each at 6.5° eccentricity. The targets were identical shapes in half the trials and different shapes in the other half, randomly selected from the set of 16 exemplars. As in the training trials the colors of the two targets, in both “same” and “different” trials, always differed by a hue angle of 60°.

**Fig 2.**
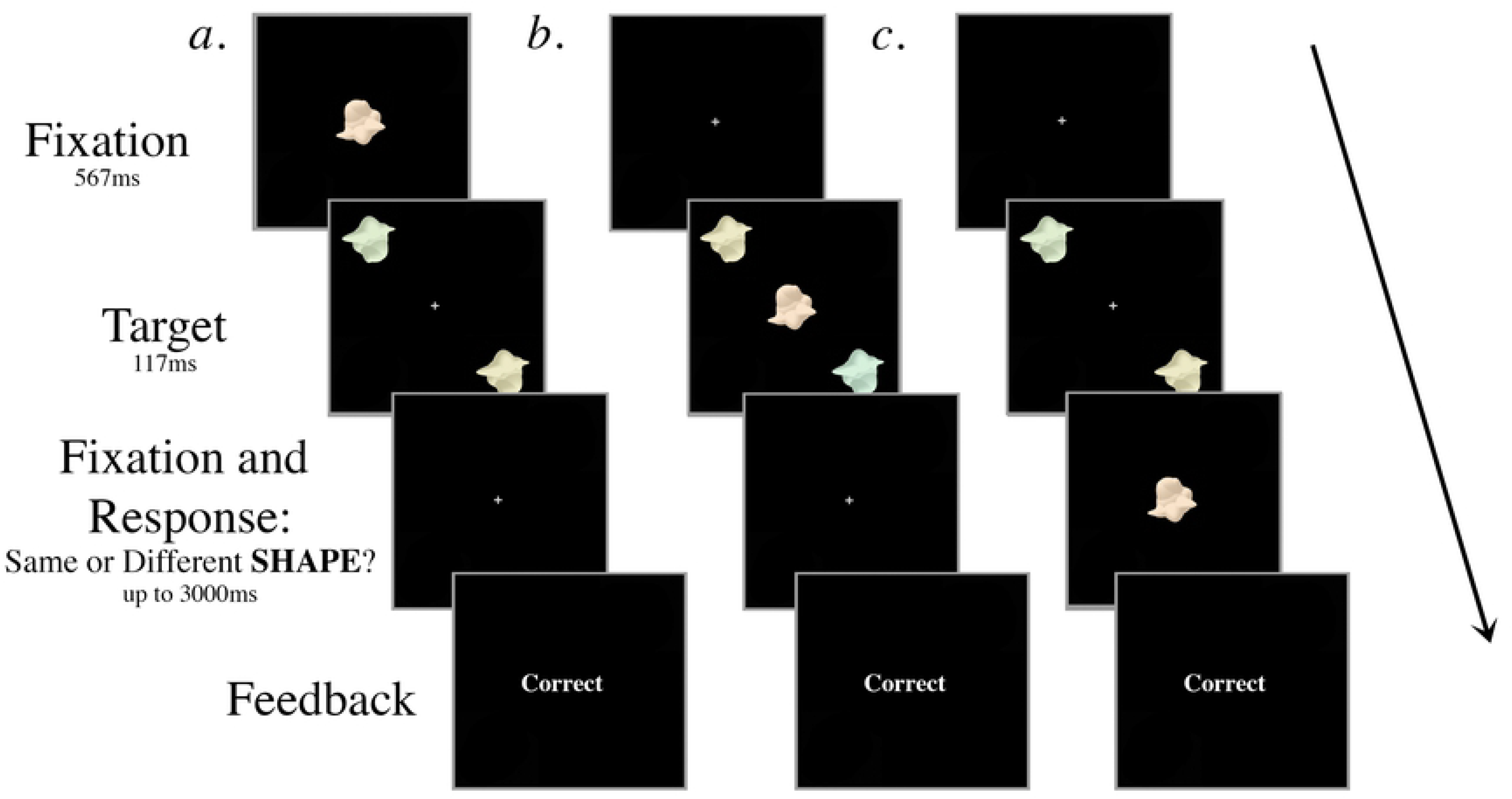
Schematic of 3 example trials in Experiment 1 with a colored distractor. Participants judged whether the peripheral targets were the same shape, ignoring their colors, which were always different. The targets and the distractor were displayed for 117ms regardless of SOA. The central distractor appeared either (a) 267ms or 117ms prior to target onset, (b) simultaneously with target onset, or (c) 117ms or 267ms after target onset. In the examples shown, the targets are different colors but identical shapes and the correct response is “same”. (In Experiment 2, participants judged whether the targets were the same color, ignoring their shapes; for these displays in Experiment 2 the correct response would be “different”.)

At one point in each trial, a distractor object appeared at fixation for 117ms. There were ten trial conditions that dictated the timing and the type of the distractor presented. First, the onset of the distractor object occurred at one of five possible SOAs: 267ms prior to target onset (−267ms), 117ms prior to target onset (−117ms), simultaneously with target onset (0ms), 117ms after target onset (+117ms), or 267ms after target onset (+267ms). Second, the distractor was either greyscale or colored with a hue angle of 63°, a color that did not occur in any of the targets. There were 80 trials for each of the ten conditions (40 “same”, 40 “different”) for a total of 800 trials in a session. All of the trial types were randomly intermingled, fully crossed, and blocked so that participants would have a chance to rest every 100 trials.

Participants were given 3s to respond after the completion of the trial before the next trial automatically commenced. As in the training task, participants used their right index finger to indicate a “same” judgment or their right middle finger to indicate a “different” judgment. Following each response, participants were given onscreen accuracy feedback.

As in training, participants were instructed to maintain fixation on the central cross throughout each trial and respond as quickly and accurately as possible. The eye-tracker was unavailable for six participants. However, given the short duration of the target display as well as the disparate peripheral target locations, any eye-movements towards the peripheral stimuli are likely to have impaired behavioral performance on the task, as only a single target would be able to be fixated (if that) during the display, which would make the second target further from fixation, making it more difficult to compare the two stimuli. In the cases of eye-tracked participants, we had to discard only 0.08% of completed trials from analysis due to eye-movements. Participants were able to complete the experimental task in ~45 minutes.

We did not include a non-distractor condition in the main experiment because the training task was effectively the discrimination task without a distractor. Additionally, a non-distractor condition differs from the experimental distractor-present condition. Thus, a ‘no-distractor’ condition would not be a good baseline as performance could be better due simply to practice or the other changes. Instead, we used performance in the −267ms SOA condition as a baseline for comparison as it is matched the experimental conditions in all key aspects with the only difference being the onset time of the distractor.

### Results

Our dependent variable was *d’* as a measure of discrimination sensitivity for comparing the targets. The hit rate was defined as the proportion of correct “same” responses on “same” trials, and the false alarm rate was defined as the proportion of “same” responses on “different” trials (see Table in S1 Table). We ran a two-way repeated measures ANOVA on *d’* with the factors of SOA (−267ms, −117ms, 0ms, +117ms, +267ms) and distractor type (grey, colored). We applied a Greenhouse-Geisser correction to the main effect of SOA in order to correct for violated sphericity found using Mauchly’s Test of Sphericity (χ^2^(9) = 21.215, *p* = 0.012). There was a significant main effect of SOA (*F*(2.75, 49.56) = 20.258, *p* < 0.001, η_p_^2^ = 0.530), no main effect of distractor type (*F*(1, 18) = 0.042, *p* = 0.841, η_p_^2^ = 0.002), and no interaction (*F*(4, 72) < 1, *p* = 0.970, η_p_^2^ = 0.007; Fig. 3). This result demonstrates that discrimination sensitivity varies with SOA, and whether the distractor object was colored or greyscale has little effect on the participants’ ability to discriminate between peripheral colored objects.

**Fig 3.**
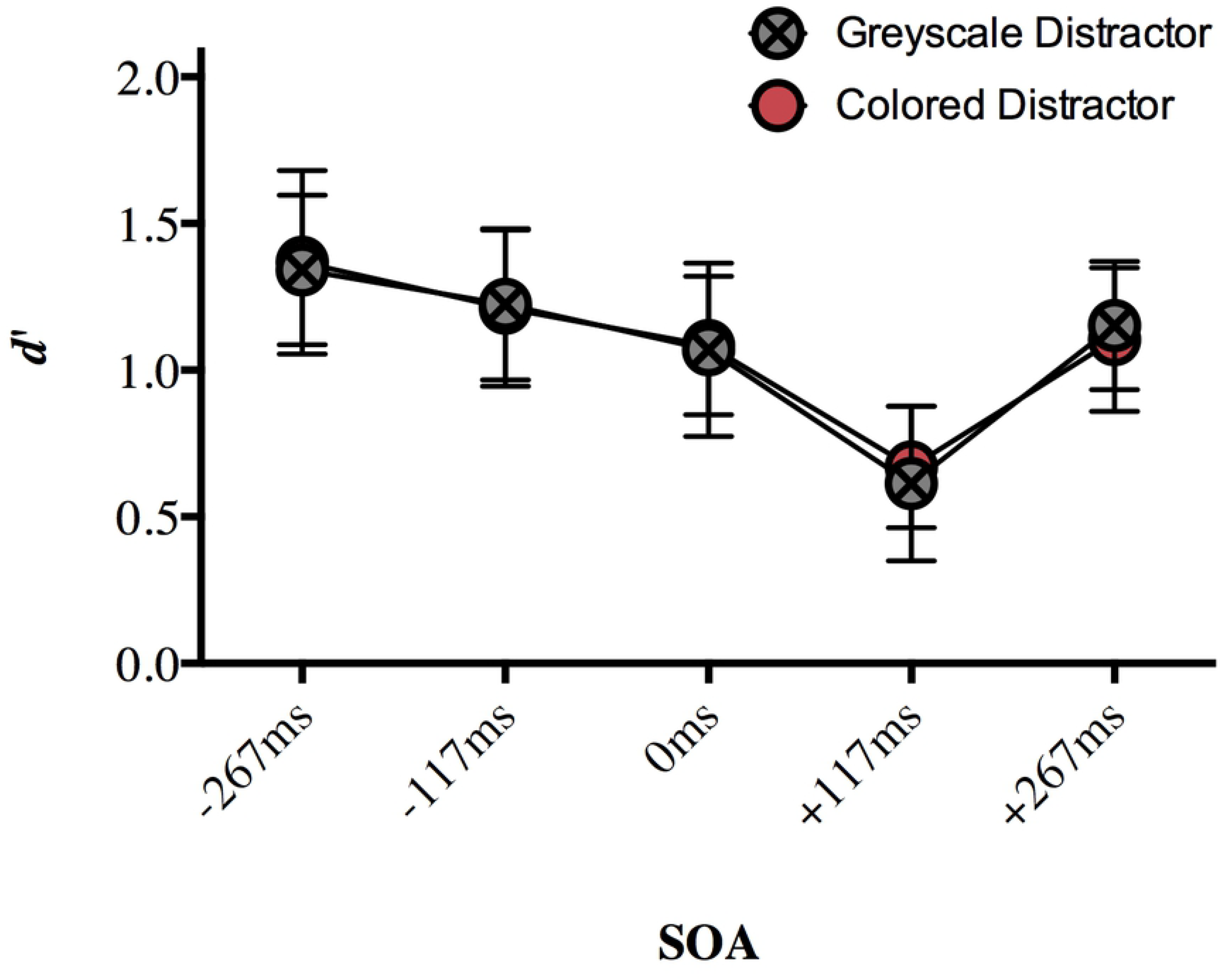
The effect of a central distractor on peripheral color discrimination (mean d’) in Experiment 1. A distractor appearing 117ms after target onset disrupted target discrimination sensitivity more than distractors appearing at every other SOA. Error bars represent 95% confidence intervals. Significant differences are discussed in text.

A Bonferroni correction for multiple comparisons (α = 0.05/10 = 0.005) was applied to post hoc analyses following up the main effect of SOA (data collapsed over distractor type). For our key SOA of +117ms, discrimination sensitivity (*d*’) was impaired compared to our relative baseline −267ms (*p* < 0.001), as well as compared to −117ms (*p* < 0.001), 0ms SOA (*p* < 0.001) and +267ms SOA (*p* < 0.001; Fig. 3). The only other significant difference was that discrimination sensitivity was significantly lower at 0ms SOA than −267ms SOA (*p* < 0.001). No other comparisons approached significance after correction (*p* > 0.005; see Table in S2 Table). Taken together, these results show that a central distractor appearing 117ms after target onset disrupted participants’ ability to discriminate between the peripheral targets more than a distractor appearing at other SOAs. This is an important replication of the foveal distractor effect [10] with stimuli that have different features.

## Experiment 2: Discriminating color

Most studies investigating the temporally-specific disruption of peripheral discrimination sensitivity have used a task requiring discrimination of fine spatial details [8,10,11, but see 9] The aim of Experiment 2 was to determine whether this foveal distractor effect would occur when participants attend to and perform a discrimination task on an object characteristic other than shape, in this case, color. Color is an object characteristic that is easily manipulated while avoiding changes to spatial details of the visual stimuli. We used the stimuli from Experiment 1 in order to minimize differences between the two experiments.

### Materials and methods

#### Participants

A naïve group of 20 participants was recruited for Experiment 2. One participant’s dataset was discarded due to chance-level performance, leaving 19 full datasets for analysis (15 female, 4 male; mean age = 21.34 ± 5.06 years). Participants reported normal or corrected-to-normal visual acuity, were screened for normal color vision using Ishihara color plates, and gave informed consent. Each received course credit or $15 for their participation.

#### Procedure

The stimuli and apparatus were the same as in Experiment 1. Prior to taking part in the experiment, participants were trained on a basic discrimination task similar to the training for Experiment 1, except that in Experiment 2, participants discriminated between the colors rather than the shapes of the objects. The shapes of the target objects in Experiment 2 were always different, randomly chosen from the set of 16 exemplars. Participants were instructed to ignore the shapes of the targets and make a judgement on whether the colors of the targets were identical or different. In each trial, one color was chosen at random between the hue angles of 0° and 200°. In “same” trials, the objects’ colors were identical. In “different” trials, the second target’s color always differed by a hue angle of 60°. The degree of difference was determined based on pilot data such that participants would be able to discriminate between the two colors with a similar accuracy as when doing the shape task described in Experiment 1, and the range of colors was chosen to complement the variability in the shapes of the exemplars. The parameters of the training task were the same as in Experiment 1 (see Fig. 1). Participants were trained until they were able to make at least 70% correct discriminations when the target array was displayed for 117ms, while maintaining fixation throughout the block.

Experiment 2 was carried out in a similar way to Experiment 1 (see Fig. 2), but the required task was different: participants were asked to judge whether the two colored objects were the same *color* while ignoring their shapes.

The eye-tracker was unavailable for seven of the participants in Experiment 2. In the cases of eye-tracked participants, we discarded 0.08% of completed trials from analysis.

### Results

Our dependent variable was again *d’* for target discrimination sensitivity. The hit and false alarm rates (see Table in S3 Table) were defined as in Experiment 1. We ran a two-way repeated measures ANOVA on *d’* with the factors of SOA (−267ms, −117ms, 0ms, +117ms, +267ms) and distractor type (grey, colored). There was a significant main effect of SOA (*F*(4, 72) = 7.328, *p* < 0.001, η_p_^2^ = 0.289), no main effect of distractor type (*F*(1, 18) = 1.045, *p* = 0.32, η_p_^2^ = 0.55), and no interaction (*F*(4, 72) = 1.918, *p* = 0.117, η_p_^2^ = 0.096; Fig. 4). This result suggests that target discrimination sensitivity on the color task varied with distractor SOA, and whether the distractor object was colored or grey had little effect on performance.

**Fig 4.**
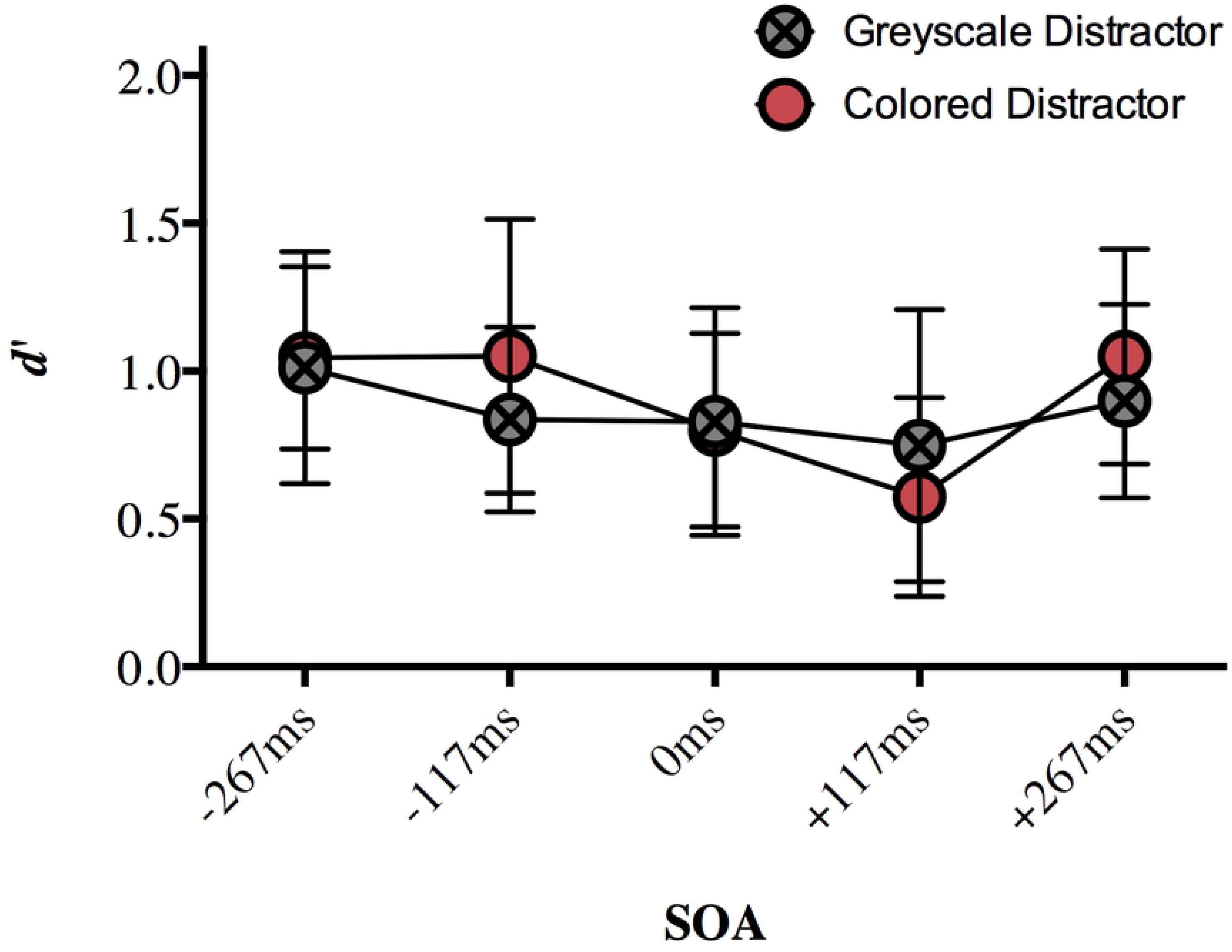
The effect of a central distractor on peripheral color discrimination (mean d’) in Experiment 2. Error bars represent 95% confidence intervals. Significant differences are discussed in text.

A Bonferroni correction for multiple comparisons (α = 0.05/10 = 0.005) was applied to post hoc analyses following up the main effect of SOA (data collapsed over distractor type). Target discrimination sensitivity was significantly impaired for +117ms SOA compared to that at −267ms SOA (*p* = 0.001), −117ms SOA (*p* = 0.003), and +267ms SOA (*p* < 0.001). No other comparisons survived correction (*p* > 0.021; see Table in S4 Table). Although the pattern is less clear for this experiment, these significant results are similar to the pattern of results from Experiment 1, where a central distractor appearing 117ms after target onset disrupted participants’ ability to discriminate between the peripheral targets more than a target appearing at other non-simultaneous SOAs. The main discrepancy is the lack of a difference between 0ms SOA and +117ms SOA, which does not come out in this experiment; being a null effect, we will not interpret this further.

## Experiment 3: Color discrimination with simple shapes

In Experiment 2, participants discriminated between the colors of novel objects. A distractor object in central vision at 117ms post-stimulus onset impaired target discrimination sensitivity relative to most of the other SOAs (except 0ms). This result suggests that feedback to foveal retinotopic cortex carries task-relevant information (in this case, color). However, the targets in Experiment 2, being novel objects, still contained complex shape information. It is therefore possible that it is not the relevance of color that drove the result, *per se*, but instead the link between the shape and color [13,14,18]. The aim of Experiment 3 was to see whether the effect at the critical SOA remained when the stimuli were restricted to a single low-level feature (color) and participants therefore did not have to ignore any aspect of the targets.

### Materials and methods

#### Participants

A naïve group of 20 participants was recruited for Experiment 3 (11 female, 9 male; mean age = 21.5 ± 3.99 years). Participants received either course credit or $15 for their participation. All participants were screened for normal color vision, and normal or corrected-to-normal visual acuity and gave informed consent.

#### Stimuli and apparatus

All aspects of the apparatus were the same as Experiments 1 and 2. The stimuli were a set of color patches, presented on a black background, using the same luminance (85), chroma (38), and hue values (0°-200°) from Experiments 1 and 2. Using Matlab, the original circles (*r* = 125 pixels) were filtered with a rotationally symmetric Gaussian low-pass filter of size 100 x 100 with a standard deviation of 10 (Fig. 5). In the experiment, the targets were sized to subtend ~1.5° visual angle as in the previous experiments.

**Fig 5.**
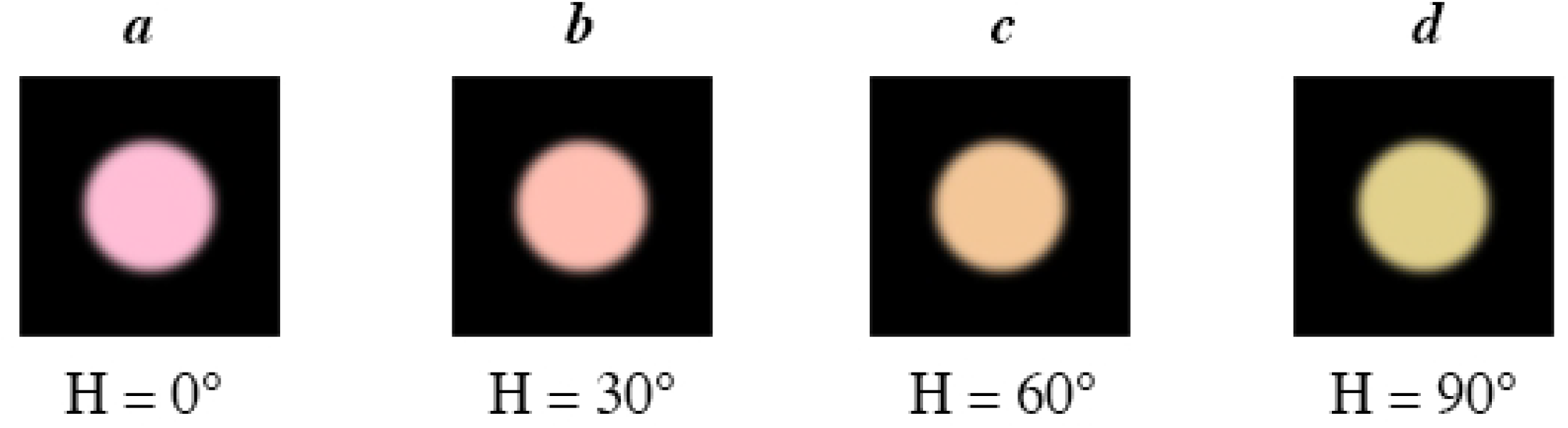
Examples of stimuli used as targets in Experiment 3. Exemplars differed by 60°, so that, for example, (a) and (b), (b) and (c), or (c) and (d) could be used as pairs. Only hue angle varied; luminance and saturation remained constant.

#### Procedure

In Experiment 3, participants were asked to judge whether the two target circles were the same or different colors. This meant that unlike in the previous experiments, they were no longer required to ignore any feature of the targets. Otherwise, the training and experimental procedures were the same as in Experiment 2. Three participants were not eye-tracked due to technical problems with the eye-tracker. For the other participants, we discarded 0.06% of the eye-tracked trials from the analysis due to fixation failures.

### Results

The dependent variable was again *d’* for target discrimination sensitivity. The hit and false alarm rates (see Table in S5 Table) were defined as in Experiments 1 and 2. A two-way repeated measures ANOVA on *d’* with the factors of SOA (−267ms, −117ms, 0ms, +117ms, +267ms) and distractor type (greyscale, colored) showed a significant main effect of SOA (*F*(4, 76) = 4.373, *p* = 0.003, η_p_^2^ = 0.187), no effect of distractor type (*F*(1, 19) = 0.117, *p* = 0.736, η_p_^2^ = 0.006), and a significant interaction (*F*(4, 76) = 4.075, *p* = 0.005, η_p_^2^ = 0.177; Fig. 6).

**Fig 6.**
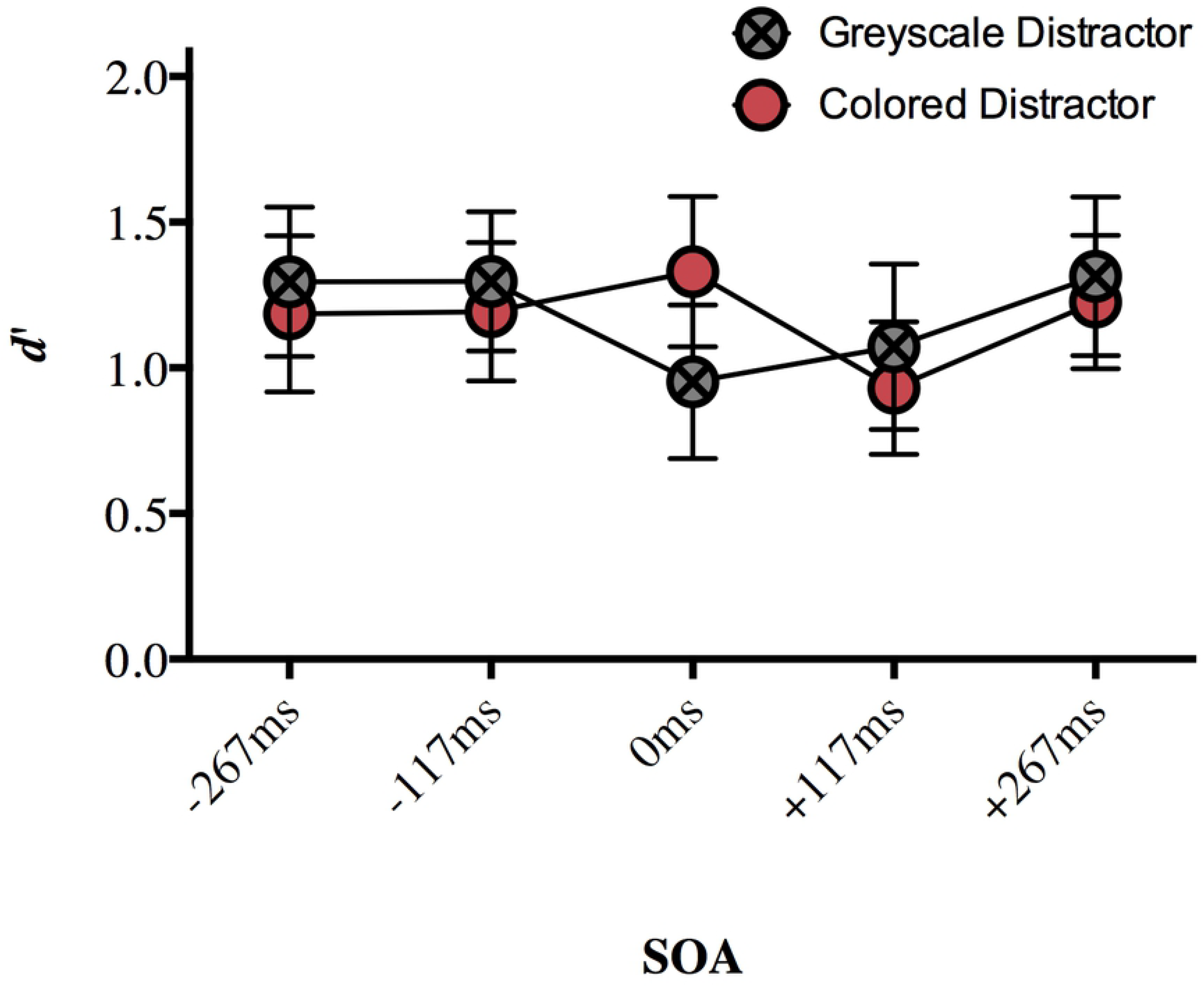
The effect of a central distractor on peripheral color discrimination (mean d’) in Experiment 3. Error bars represent 95% confidence intervals. Significant differences are discussed in text.

We followed up the interaction with a repeated measures ANOVA on the distractor type conditions separately (Fig. 6). There was a main effect of SOA for both the colored distractor (*F*(4, 72) = 9.659, *p* < 0.001, η_p_^2^ = 0.349) and greyscale distractor (*F*(4, 72) = 13.026, *p* < 0.001, η_p_^2^ = 0.42; Fig. 6) conditions. A Bonferroni correction for multiple comparisons (α = 0.05/20 = 0.0025) was applied to the post hoc analyses. For the colored distractor condition, target discrimination sensitivity was impaired with the distractor was presented at +117ms SOA compared to a distractor presented at 0ms SOA (*p* = 0.002; Fig. 6). The difference in mean *d’* values for +117ms and −267ms (*p* = 0.058), −117ms (*p* = 0.013), and +267ms (*p* = 0.003) did not reach significance after correction but suggest a pattern of results similar to that demonstrated in Experiments 1 and 2 (Table in S6 Table). No other comparisons approached significance (*p* > 0.05).

For the greyscale condition, discrimination sensitivity was significantly impaired for 0ms SOA compared to −267ms SOA (*p* = 0.001). Mean *d’* values for +117ms SOA were numerically lower than mean *d’* values at −267ms SOA (*p* = 0.019), −117ms SOA (*p* = 0.007), and +267ms SOA (*p* = 0.019) but these differences did not reach significance after correction (Fig. 6).

We also followed up the interaction of SOA and distractor type by examining the effect of distractor type at each SOA separately. Post-hoc analyses of the interaction using a Bonferroni correction (α = 0.05/10 = 0.005) showed that distractor color affected discrimination accuracy when presented simultaneously with the targets, with a grey distractor impairing target discrimination sensitivity relative to a colored distractor (*p* = 0.004; Fig. 6). At all other SOAs, there was no significant effect of the color of the distractor type (*p >* 0.05; Table in S7 Table).

## Discussion

The aim of this study was to test whether the foveal distractor effect is limited to form-related information or extends to other visual features. In Experiment 1 we used colored novel objects to replicate the effect first demonstrated with achromatic stimuli in Weldon et al. [10]. When participants were asked to discriminate two peripheral target objects on shape while ignoring their color, a distractor presented at fixation at +117ms SOA impaired perceptual discrimination more than a distractor presented at SOAs very early or late in the trial. At the critical SOA, the targets are no longer present onscreen and the distractor appears in an entirely different location from that of the target array.

We followed up Experiment 1 by asking whether we would see the same pattern in the data if participants were asked to discriminate color, rather than shape, on the same set of stimuli. We demonstrated the foveal distractor effect in Experiment 2, where participants were required to ignore the targets’ shapes (which were always different), and discriminate object color. Although the results were not as clear as in Experiment 1, this is the first demonstration of the foveal distractor effect during a color discrimination task. Although this result might indicate that feedback to foveal retinotopic cortex carries task-relevant information that is not limited to object shape (in this case, color), our targets in Experiment 2 still contained complex shape information. It is possible that the feedback of bound color-shape information rather than the feedback of color in and of itself might be driving the behavioral effect. We addressed this in Experiment 3 by minimizing shape information in the stimuli and presenting homogenous color patches as targets.

In Experiment 3, we found that the delayed disruption of peripheral discrimination sensitivity to be somewhat diminished. This result is consistent with evidence that foveal feedback is selectively employed for tasks that involve spatial detail [9]. Foveal vision, in contrast with peripheral vision, is highly specialized and ideal for tasks involving stimuli with fine spatial details. One explanation of the function of a foveal feedback mechanism is that foveal cortex acts as a high-resolution buffer [19,20], or visual “scratchpad”, for storing task-relevant information [6,9]. This specialized region of cortex may be recruited for the purpose of resolving perceptual decisions during difficult perceptual tasks with peripheral stimuli. If this is the case, foveal feedback would be less engaged for a task that requires discrimination of objects with fewer spatial details (as in Experiment 3), even when the task is similarly challenging for observers. Our behavioral result here supports this explanation, but converging evidence from fMRI studies that take advantage of MVPA techniques is necessary to compare foveal retinotopic cortex content during different types of tasks.

Experiment 3 also had an interaction of distractor type and SOA, unlike in Experiment 1 and 2 where we found only a main effect of target SOA. A greyscale distractor was more disruptive to discrimination sensitivity than a colored distractor when it was presented simultaneously with the targets (and only then), which suggests the disruption is a result of a grey distractor causing some differential interference with *feedforward* processing (as opposed to feedback) of color stimuli. Although central distractors that are irrelevant to targets have been shown to cause more interference in visual search tasks than when the central distractor is identical to the target [21], this finding differs somewhat from previous work where participants performed a discrimination task on achromatic versions of the stimuli used in this paper. A central, inconsistent distractor (an angular, “cubie” version of the target stimuli, [7]) did not interfere with peripheral discrimination sensitivity more than a distractor that was consistent with the targets (an object from the same shape category) [10]. The different effect is perhaps driven by computational differences related to the chromaticity of the distractor [22], the requirement to discriminate between a feature with low spatial frequency rather than complex spatial information [9], or some combination of these possibilities. Although this finding is intriguing, any distractor appearing on the display simultaneously with the distractors could reasonably be expected to interfere with peripheral perception simply because there is more information present in the visual field. Furthermore, this finding at 0ms SOA is not directly relevant to our main investigation regarding feedback at later SOAs, so we will not speculate more here.

The timecourse of the effect described in this paper for this particular paradigm has been consistent across multiple experiments (see 5). The ~40ms discrepancy between the timecourse demonstrated here and Yu and Shim [11], though they target a similar effect, may be due to the difference in perceptual task. Fan et al. [9] showed that foveal interference occurs around 450ms SOA for trials involving an additional mental rotation task. The difficulty of a task and/or specific task demands may determine the time at which feedback to the foveal cortex occurs, and thus, the time at which our foveal distractor effect would be evident.

Another possible explanation for these discrepancies may be due to the differences in temporal relationship between the offset of the target and the onset of the distractor on the visual display. In the studies presented here, the *offset* of the target array coincides with the *onset* of the distractor at the critical SOA of +117ms. Although there is no overlap between the display time of the target array and the display time of the distractor at 117ms SOA, we cannot be sure based on the limited amount of literature using this paradigm. The behavioral paradigm designed to target this effect is still new; further experiments are necessary to map out the timecourse of the effects reported here and elsewhere [9–11].

Overall, the present experiments demonstrate that the foveal distractor effect is not specific to object shape information, but that feedback to the foveal confluence is also important for the peripheral discrimination of color, especially when discriminating between <u>colored complex</u> shapes. The more subtle effect of a central distractor at +117ms during discrimination of homogenous color patches suggests that the foveal feedback signal is flexible and may be related to tasks involving discrimination between fine spatial detail [9]. In Williams et al. [6], object information in foveal cortex was present only when participants performed the object discrimination task, but not when they performed a color discrimination task on the same stimuli. It will be important to employ neuroimaging studies to determine whether color information is likewise present at foveal retinotopic cortex only during a behavioral task requiring color discrimination. Such studies would also be able to address the question of whether irrelevant information is fed back to foveal cortex (which is more difficult to measure behaviorally). That said, the evidence from the present set of experiments, namely, that perception of peripheral object form and color is affected by disruption of foveal representations at a timepoint consistent with feedback to foveal cortex, lends credence to the proposal that foveal retinotopic cortex serves to store or compute task-relevant visual information during difficult perceptual tasks.

